# Trimester-specific Zika virus infection affects placental responses in women

**DOI:** 10.1101/727081

**Authors:** Fok-Moon Lum, Vipin Narang, Susan Hue, Jie Chen, Naomi McGovern, Ravisankar Rajarethinam, Jeslin J.L. Tan, Siti Naqiah Amrun, Yi-Hao Chan, Cheryl Y.P. Lee, Tze-Kwang Chua, Wearn-Xin Yee, Nicholas K.W. Yeo, Thiam-Chye Tan, Xuan Liu, Sam Haldenby, Yee-sin Leo, Florent Ginhoux, Jerry K.Y. Chan, Julian Hiscox, Chia-Yin Chong, Lisa F.P. Ng

## Abstract

Zika virus (ZIKV) infection during pregnancy is associated with neurologic birth defects, but the effects on placental development are unclear. Full-term placentas from three women, each infected with ZIKV during specific pregnancy trimesters, were harvested for anatomic, immunologic and transcriptomic analysis. In this study, each woman exhibited a unique immune response, but they collectively diverged from healthy controls with raised IL-1RA, IP-10, EGF and RANTES expression, and neutrophil numbers during the acute infection phase. Although ZIKV NS3 antigens co-localized to placental Hofbauer cells, the placentas showed no anatomical defects. Transcriptomic analysis of samples from the placentas revealed that infection during trimester 1 caused a disparate cellular response centered on differential eIF2 signaling, mitochondrial dysfunction and oxidative phosphorylation. These findings should translate to improve clinical prenatal screening procedures for virus-infected pregnant patients.

## Background

Zika virus (ZIKV) caused numerous outbreaks of infection worldwide [1] and the scale of these outbreaks highlighted several features of ZIKV infection that had previously been unrecognized or under reported. One such feature included congenital fetal growth-associated anomalies as a result of ZIKV infection during pregnancy, termed Congenital Zika Syndrome (CZS) [2, 3]. Pregnancy is divided into three trimesters, based on the series of developmental changes that occur in the fetus and the physiological changes that occur in the mother. Understanding the nature of the ZIKV–host interactions at each trimester is required to determine the consequences of maternal–fetal ZIKV transmission and its impact on CZS development. Important physiological changes occur in women during pregnancy, including immune responses [4]. The unique immunological state experienced during pregnancy prevents the mother from rejecting the fetus [5], but also increases susceptibility to infection [6].

The placenta develops from trophectoderm surrounding the blastocyst and rapidly grows after implantation to establish the fetal life support system before the rapid growth of the fetus in the second half of pregnancy. The placenta is the physical barrier between the mother and her fetus. It is the site of transport of oxygen nutrients, antibodies and waste products between mother and baby [7]. Importantly, the placenta also protects the fetus from infections, exhibiting a robust innate and adaptive immune response to pathogens [8]. Hofbauer cells, are macrophages located within the stroma of the placenta and can be isolated from the placenta and be infected by ZIKV experimentally [9]. Positive ZIKV infection has been detected in placental tissue [10] and fetuses [3] of women who have terminated their pregnancies, suggesting the virus can evade placental immune defense mechanisms. As the risk of congenital anomalies, including microcephaly, is largely associated with ZIKV infection during the first trimester of pregnancy [11], it is also possible that ZIKV infection perturbs placental development. Such perturbations could also upset the placental immune function. The mechanisms underlying how ZIKV infection in the three trimesters affects the placental cellular responses, especially in pregnancies without any congenital complications, are currently unknown. Understanding these provide an alternate perspective regarding ZIKV pathogenesis in asymptomatic pregnant patients.

Here, the placental cellular responses in ZIKV-infected pregnant women were probed and the study delineated how ZIKV infection affected placental development in each trimester. Three pregnant women were recruited to the study who had been infected with ZIKV during the first, second or third trimesters. After successful, full-term delivery of healthy infants, the placentas were harvested and separated into the placental discs (chorionic villi) and fetal membranes (chorion and amnion) to investigate the cellular responses in these anatomically distinct structures. Histology and immunofluorescent analyses were used to identify any tissue parenchymal abnormalities and immune-cell changes. This was combined with RNA-sequencing (RNA-seq) to determine the transcriptomic profiles of these samples. Although the number of ZIKV-infected pregnant women recruited was low and may not represent the entire population, it still provides insightful data on the possible placental responses during trimester-specific ZIKV infection.

## Methods

### Ethics approval and consent to participate

Written informed consent was obtained from all participants in accordance with the Declaration of Helsinki for human research. The study protocol was approved by the SingHealth Centralized Institutional Review Board (CIRB Ref: 2016/2219). Blood samples were collected from healthy donors who had provided written consent, in accordance with the guidelines of the Health Sciences Authority of Singapore (study approval number: NUS IRB: 10-250).

### Patients and sample collection

Three pregnant women were enrolled into the study upon admission to the KK Women’s and Children’s Hospital, Singapore. Upon admission, patients were confirmed to be infected via qRT-PCR detection of ZIKV RNA in their urine. Whole blood samples were collected at two time points: 0-7 days post illness onset (PIO) and 10-14 days PIO, representing the acute and convalescent phases, respectively. Whole blood was collected in EDTA Vacutainer tubes (Becton Dickinson) after peripheral venipuncture. Each sample was first used for blood count, whole blood staining and viral load quantification and then centrifuged at 256 g for 10 min to collect plasma for storage at −80°C. Whole blood samples were also collected from three healthy women and pre-screened for ZIKV viral RNA and ZIKV-specific antibodies using in-house protocols [12]. Placentas from two healthy individuals were obtained as control. All healthy donors were non-febrile and had no signs of acute illness during recruitment.

### Placenta processing

After successful full-term delivery, the placenta was maintained on ice and transported to the Singapore Immunology Network for immediate downstream processing within 2 h of expulsion. The term placenta was first separated into the fetal membrane (chorion and amnion) and placenta disc (chorionic vili) layers. A small section of the tissue was removed and placed in 10% neutral buffered formalin (NBF; Sigma Aldrich) for histology and immunofluorescence staining. The remaining tissues were finely diced in a digestion medium containing collagenase type 4 (0.2mg/ml) and DNAse (0.05mg/ml) prepared in Roswell Park Memorial Institute medium (Hyclone) containing 10% Fetal Bovine Serum (Gibco). Refer to supplemental materials for more information.

The fetal membranes were incubated in digestion medium for 3 h in a 37°C incubator supplemented with 5% CO_2_. The fetal cells were then centrifuged (400 g, 5 min) and repeatedly washed with PBS to remove traces of digestion medium before red blood cell (RBC) lysis. Lysis was done with in-house RBC lysing solution containing 155 mM NH_4_C1 (Sigma Aldrich), 10 mM KHCO_3_ (Sigma Aldrich) and 0.1 mM EDTA (Life Tech). Lysed cells were then enumerated and used for downstream procedures.

The placenta disc were incubated in digestion medium for 1 h, and then the digestion medium was carefully overlaid on Ficoll-Paque Plus (GE Healthcare) at a ratio of 2:1, respectively. The samples were then centrifuged at 1,600 g for 20 min (with minimal acceleration and deceleration) to isolate the “buffy coat” containing leucocytes. The leucocytes were carefully removed and traces of RBCs were subsequently lysed. The cells were enumerated and used for downstream procedures.

In both cases, the digestion medium was filtered through a 100μM filter unit after digestion to remove any undigested tissues. The volume of medium used was adjusted according to the size of the tissue being digested.

### Histology and immunofluorescence

Placental tissues were first fixed in 10% NBF (Sigma Aldrich) at room temperature for 24 h before being processed for routine histologic evaluation. Briefly, isolated tissues were embedded in paraffin wax, cut into 5 μM thick sections, deparaffinized and then stained with Hematoxylin and Eosin (H&E). The stained sections were viewed under an Olympus BX53 upright microscope (Olympus Life Science, Japan), and images were captured with an Olympus DP71 digital camera using an Olympus DP controller and DP manager software. All images were evaluated by a certified pathologist.

Tissue sections were stained by standard immunofluorescence technique. In brief, antigen-retrieval was performed after deparaffinisation, using DAKO target retrieval solution, pH9. After washing with TBS-T, the tissues were then treated with 10% goat serum for 30 min to block the nonspecific reaction. The tissues were then incubated overnight with various combinations of a ZIKV in-house antibody [13] (1:100) and CD 163 (Thermofisher; 1:50) at 4°C. The slides were incubated with secondary antibodies (1:500) such as Alexa Fluor^®^ 488 goat anti rabbit IgG (Life technologies) and Alexa Fluor^®^ 594 goat anti mouse IgG (Invitrogen) for 30 min in the dark, and the nuclei were counter stained with Vectashield™ Hard Set mounting medium with DAPI. The stained slides were examined under Nikon eclipse 90i fluorescence microscope (Nikon, Japan) and images were captured with microscopic camera, DS-Fi3 (Nikon, Japan) using NIS Elements Imaging software, (Nikon, Japan).

### Blood count

A complete blood count (CBC) was determined using an Ac·T diff hematology analyser (Beckman Coulter), according to manufacturer’s instructions. Beckman Coulter 4C^©^ Plus Tri-Pack Cell Controls (Beckman Coulter) were used to confirm instrument accuracy and precision.

### Whole blood labeling and flow cytometry

Whole blood (100 μL) was labeled as previously described [13]. Antibodies were used to identify CD45+ leucocytes (mouse anti-human CD45, Biolegend), high SSC-A CD16+ neutrophils (mouse anti-human CD16, Biolegend), CD14+ monocytes (mouse anti-human CD14, BD Biosciences), CD3+ T cells (mouse anti-human CD3, BD Biosciences), CD4+ T helper cells (mouse anti-human CD4, eBioscience), CD56+ NK cells (mouse anti-human CD56, Miltenyi Biotec) and CD19+ B cells (mouse anti-human CD19, eBioscience). Cell fixation and RBC lysis was performed using 1X FACS lysing solution (BD Biosciences), and permeabilization using 1X FACS permeabilization solution 2 (BD Biosciences) before staining with a ZIKV NS3 protein-specific rabbit polyclonal antibody [13]. The labeled cells were counter-stained with a fluorophore-tagged secondary goat anti-rabbit IgG (H+L) antibody (Invitrogen), before acquisition on a LSR Fortessa (BD Biosciences). Dead cells were excluded using a Live/Dead^TM^ Fixable Aqua Dead Cell Stain Kit (ThermoFisher Scientific). The frequencies of peripheral blood immune subsets were obtained with the following formula: Percentage of specific immune subset (obtained from immune-phenotyping) × total leucocyte number (obtained from CBC) = cellular number of specific immune subset.

### Multiplex microbead immunoassay for cytokine quantification

Cytokine and chemokine levels in plasma from ZIKV-infected patients were measured simultaneously using a multiplex microbead-based immunoassay, ProcartaPlex Human Cytokine/Chemokine/Growth Factor Panel 1 (Thermo Scientific), as previously described [14]. Plasma samples and reagents were prepared, and immunoassay procedures were performed according to the manufacturers’ instructions.

### ZIKV virion-based ELISA

The presence and titers of ZIKV-specific antibodies in the plasma of the infected pregnant women at both acute and convalescent phases were determined using a ZIKV virion-based ELISA, as previously described [12]. Plasma samples were tested at 1:200 dilutions.

### RNA sequencing (RNA-seq)

RNA samples, extracted from the fractions of isolated placental cells, were treated with DNase using an Ambion Turbo DNA-free Kit (Ambion), and then purified using Ampure XP beads (Agencourt). The DNase-treated RNA (2 μg) was treated with Ribo-Zero using an Epicentre Ribo-Zero Gold Kit (Human/Rat/Mouse) (Epicentre) and re-purified on Ampure XP beads.

Successful RNA depletion was verified using a Qubit (Thermo Fisher Scientific) and an Agilent 2100 Bioanalyzer (Agilent) and all depleted RNA was used as input material for the ScriptSeq v2 RNA-Seq Library Preparation protocol. RNA was amplified for 14 cycles and the libraries were purified on Ampure XP beads. Each library was quantified using a Qubit and the size distribution was assessed using an AATI Fragment Analyser (Advanced Analytical); the final libraries were pooled in equimolar amounts. The quantity and quality of each pool was assessed using a Fragment Analyser and by qPCR using an Illumina Library Quantification Kit (KAPA Biosystems) on a Light Cycler LC480II (Roche), according to manufacturer’s instructions. The template DNA was denatured according to the protocol described in the Illumina cBot User guide and loaded at a 12 pM concentration. Sequencing was carried out on three lanes of an Illumina HiSeq 2500 with version 4 chemistry, generating 2 × 125 bp paired-end reads.

### RNA-seq data analysis

Quality assessment of the RNA sequencing reads was performed using FASTQC (version 0.11.5; http://www.bioinformatics.bbsrc.ac.uk/projects/fastqc). Cutadapt (version 1.2.1) [15] was used to remove Illumina adapter sequences with option −O 3 so that the 3’ end of any reads that matched the adapter sequence for ≥3 bp were trimmed. The reads were further trimmed using Sickle (version 1.20; https://github.com/najoshi/sickle) with a minimum window quality score of 20. Reads <10 bp after trimming were removed. The resulting clean, paired-end reads were mapped to human transcript sequences obtained from Gencode (version 27) [16] using Salmon (version 0.9.1) [17]. Transcript abundances quantified by Salmon were summarized to gene-level counts and normalized gene-level abundances in transcript per million (TPM) units using the tximport R/Bioconductor package (version 1.2.0) [18].

Principal component analysis (PCA) was performed on all samples using genes with TPM > 1 in at least one sample. The TPM values were log_2_ transformed after adding a pseudocount of 1.0 to prevent negative values, and used for PCA in FactoMiner R package (version 1.33) [19] with default scaling transformation.

Differential gene expression (DGE) analysis was performed using edgeR R/Bioconductor package (version 3.16.5) [20]. Gene-level counts for samples relevant to the conditions being compared were loaded into edgeR and a negative binomial (NB) generalized linear model (GLM) extended with quasi-likelihood (QL) methods was fitted to the counts data. As recommended for gene-level counts summarized using tximport, the transcript abundances and average transcript lengths of genes were used to specify the offset term in the edgeR model in order to correct for potential differences in transcript usage between samples [18]. Low expressed genes with <5 counts in all samples were removed. The trended NB dispersion was estimated using the estimateDisp function and the QL dispersion was estimated using the glmQLFit function. Differential gene expression was determined using a quasi-likelihood F-test (glmQLFTest function), which provides a conservative and rigorous type I error rate control [20]. Genes with a Benjamini-Hochberg multiple testing corrected p-value <0.05 were considered differentially expressed. Differentially expressed gene lists were analyzed with DAVID gene ontology analysis (version 6.8) [21], Ingenuity Pathway Analysis^®^ (IPA) and Gene Set Enrichment Analysis (GSEA) (version 3.0) [22] to identify associated biological conditions and pathways. The curated gene sets used for GSEA included the hallmark (H) and immunologic signatures (C7) gene sets downloaded from Broad Institute’s Molecular Signatures Database that accompanies the GSEA software.

### Multiple factor analysis

Data from immunophenotyping of whole blood and cytokine quantification in plasma were analyzed together using multiple factor analysis (MFA) [23]; MFA permits several groups of variables to be studied on the same set of individuals. Analysis was performed with nine individual samples, including the three ZIKV-infected patients in the acute and convalescent phases of infection, and the three healthy controls. The immunophenotyping and cytokine measurements were specified as two different groups of variables consisting of log_10_ transformed cell counts per µl of whole blood and log_10_ transformed cytokine levels in pg/ml units. The analysis was carried out using R package FactoMineR (version 1.33) [19] with the default parameters. The data were plotted using custom R scripts. Data from placenta immunophenotyping and RNA sequencing were also analyzed using MFA. Data acquired from placenta disc and fetal membrane fractions were analyzed separately. The individuals in this analysis included the three ZIKV-infected patients and two healthy controls. Cellular fractions were expressed as a percentage of the parent gate in FACS analysis and RNA-seq measurements were expressed in log_2_ TPM units.

## Results

### Pregnant women in different trimesters exhibited differential susceptibility to ZIKV infection

Three ZIKV-positive pregnant women were recruited to the study, who had been infected with ZIKV in different trimesters (Table 1). Blood samples were taken during the acute (5-6 days post-illness onset (PIO)) and convalescent (9-14 days PIO) phases of infection, and analyzed for the presence of ZIKV NS3 antigens.

**Table 1.** Profiles of ZIKV-infected pregnant patients

|  | Acute phase |  |  | Convalescent phase |  |  |  |
| --- | --- | --- | --- | --- | --- | --- | --- |
|  | Patient 1 | Patient 2 | Patient 3 | Patient 1 | Patient 2 | Patient 3 | Healthy<sup>g</sup> |
| Samples collection (PIO)<sup>a</sup> | 5 | 7 | 6 | 14 | 13 | 9 | - |
| Age (years) | 33 | 34 | 28 | - | - | - | - |
| Presence of symptom<sup>b</sup> |  |  |  |  |  |  |  |
| *Fever* | No | No | Yes | - | - | - | - |
| *Headache* | Yes | No | Yes | - | - | - | - |
| *Rash* | Yes | No | No | - | - | - | - |
| *Pain around eye area* | Yes | No | No | - | - | - | - |
| Presence of ZIKV RNA in urine<sup>c</sup> | Yes | Yes | Yes | - | - | - | - |
| Percentage ZIKV antigen in blood | 15 | 0.44 | 2.66 | 7.61 | 0.19 | 10.1 |  |
| (Infection level)<sup>d</sup> | (moderate) | (low) | (low) | (low) | (low) | (moderate) | - |
| Cell counts (numbers/μl blood)<sup>e</sup> | | | | | | | |
| *Neutrophils* | 9378 | 4915 | 4868 | 8934 | 3668 | 5936 | 3784 ± 428 |
| *Monocytes* | 243 | 257 | 37 | 323 | 239 | 99 | 253 ± 96 |
| *CD4 T cells* | 301 | 954 | 393 | 501 | 491 | 407 | 993 ± 105 |
| *CD8 T cells* | 129 | 341 | 125 | 237 | 236 | 154 | 358 ± 83 |
| *DNT cells* | 41 | 30 | 27 | 132 | 29 | 25 | 93 ± 30 |
| *NK cells* | 57 | 72 | 126 | 232 | 103 | 115 | 163 ± 75 |
| *NKT cells* | 81 | 53 | 79 | 264 | 50 | 69 | 87 ± 39 |
| *CD56hi cells* | 16 | 13 | 15 | 28 | 12 | 15 | 11 ± 1 |
| *CD14+CD56+ cells* | 30 | 43 | 9 | 41 | 43 | 29 | 31 ± 13 |
| *B cells* | 74 | 309 | 85 | 105 | 114 | 82 | 211 ± 100 |
| Antibody titer (fold increase)<sup>f</sup> |  |  |  |  |  |  |  |
| *IgM* | 1.29 | 1.91 | 2.72 | 0.92 | 1.79 | 2.68 | - |
| *IgG* | 1.10 | 3.79 | 3.14 | 0.88 | 3.85 | 2.86 | - |

<sup>a</sup> Samples were collected during both acute (1-7 days post-illness onset (PIO)) and convalescent (8-14 days PIO) phases. Patient 1, 2 and 3 were infected during the first, second and third trimester, respectively.
<sup>b</sup> Presence of any symptoms were recorded during the patients' first visit to the hospital
<sup>c</sup> Presence of ZIKV RNA in urine samples was determined by qRT-PCR during the patients' first visit to the hospital.
<sup>d</sup> Infection is determined by the percentage of ZIKV Ag positive CD14+ monocytes and the level of infection can be classified as moderate (>10%) or low (<10%)
<sup>e</sup> Cellular numbers of the peripheral blood immune subset were calculated with the following formula: [Percentages of specific immune subset × total leucocyte numbers = cellular numbers of specific immune subset]. Percentages of specific immune subset were obtained with immune-phenotyping. Total leucocyte numbers were obtained with a hematology analyzer.

CD14+ monocytes are the predominant cellular target of ZIKV [13]. Interestingly, each patient showed a different ZIKV NS3 antigen profile in CD14+ monocytes. 15% of Patient 1’s CD14+ monocytes were positive for ZIKV NS3 antigens, which persisted to a lesser degree in the convalescent phase (7.61%). In Patient 2, ZIKV NS3 antigens were barely detectable in CD14+ monocytes at both time-points (019-0.44%). Lastly, Patient 3 exhibited a “delayed” infectious response, in which 2.66% of CD14+ monocytes were positive for ZIKV NS3 antigen levels during the acute phase, which increased to 10.1% of total CD14+ monocytes during the convalescent phase (Table 1 and Supplementary Figure 1A and 1B). The different infection profiles exhibited by the three patients were corroborated by variations observed in their blood immune-cell numbers (Table 1 and Supplementary Figure 2A and 2B) and ZIKV-specific antibody titers (Table 1 and Supplementary Figure 1C and 1D). Interestingly, ZIKV-specific antibody production was weak in Patient 1, perhaps indicating a direct consequence of ZIKV-infection on the host humoral immune response during the first trimester. Titers of ZIKV-specific antibodies (both IgM and IgG) did not change during the convalescent phase in both Patients 2 and 3 (Table 1 and Supplementary Figure 1C and 1D).

### ZIKV antigen localized to placental Hofbauer cells and persisted until birth

All three patients successfully carried infants to term with no microcephaly or CZS and infants were delivered via normal vaginal deliveries with no intrapartum complications. Once delivered, the full-term placenta was separated into placental discs and fetal membranes, and a series of histological analyses were performed. By H&E staining of the placenta discs, villous maturity within the normal limits of a term placenta was observed in all three cases. The chorionic villi of all three placental discs showed patchy dilatation featuring stromal oedema and prominent aggregates of vacuolated cells. These vacuolated stromal cells have the morphologic features of villous stromal macrophages i.e. Hofbauer cells. No features of ongoing or remote acute or chronic villitis, intervillositis, villous necrosis or fibrosis, was observed (Figure 1A). This indicated that ZIKV infection did not induce any overt adverse placenta pathology. The fetal membranes of all three placentas also showed several reactive changes, including increased numbers of vacuolated mononuclear cells within the subamniotic connective tissue, in keeping with increased infiltration of macrophages. In addition, the amniotic epithelial cells exhibited columnar cell metaplasia, being more pronounced in placentas infected in the first and second trimesters. There are no signs of acute chorioamnionitis (Figure 1B).

**Figure 1.**
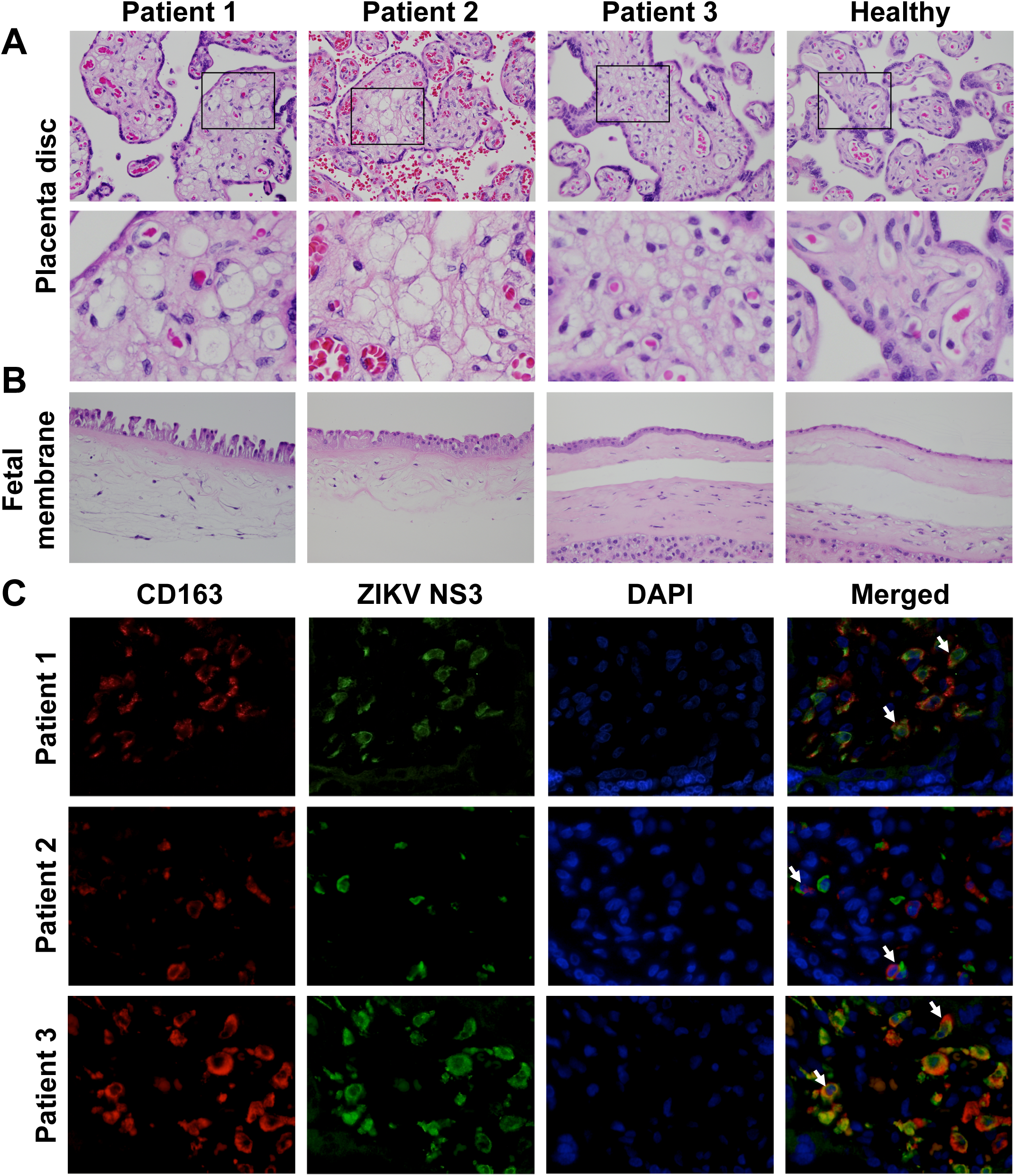
ZIKV NS3 antigen in CD163-positive macrophages impacts stromal changes in full-term placentas. Representative hematoxylin and eosin (H&E)-stained sections showing the placenta disc (chorionic villi) and fetal membranes of three patients infected by ZIKV at different gestational age. (A) Aggregates of vacuolated cells morphologically in keeping with Hofbauer cells are observed in the chorionic villi of all three infected patients. Areas highlighted by the black boxes are magnified in the lower panel. (B) Increased stromal and mononuclear cells are noted in the sub-amniotic connective tissue of the fetal membranes of the infected patients. All images were captured at 40X magnification. (C) Immunofluorescence microscopy was used to visualize ZIKV NS3 antigen (green) and CD163 protein (red) in the full-term placentas of the ZIKV-infected patients. The white arrows indicate co-localisation of the ZIKV antigen and CD163 protein within the villous stroma of the placenta disc. All images were captured at 40X magnification.

Immunofluorescence staining of the placental discs confirmed the presence of CD163+ Hofbauer cells, which exhibit functions similar to those of M2-like macrophages [24]. Counter-staining with a ZIKV NS3-specific antibody [13] indicated that ZIKV protein co-localized to Hofbauer cells (Figure 1C), in line with previous reports [9]. Staining of placental discs with secondary antibody alone, did not result in any signals (Supplementary Figure 3). Thus, this ruled out autofluoresence and indicated positive infection of the placenta, regardless of the pregnancy trimester in which ZIKV infection occurred. Notably, these data showed that ZIKV proteins were present in the placenta up to delivery, without causing any physical harm to the newborn infant. However, it is important to continue monitoring the newborn as anomalies may manifest at a later time [25].

### The ZIKV immune response correlated with the level of ZIKV infection in CD14+ monocytes

The immunological response to ZIKV infection in the three pregnant women was compared between the acute and convalescence time-points, which will provide important insights on how the host response is affected by the infection in different trimesters. Blood plasma was assayed for 42 different immune mediators, including pro-inflammatory and anti-inflammatory cytokines, chemokines and growth factors. Plasma samples from healthy, non-pregnant females were used as controls. To segregate the patients based on immunological response, the derived data was processed by multiple factor analysis (MFA) [23], which collectively analysed different measurements made from the same set of individuals. Both datasets were specified as two different groups of measurements to the algorithm.

The unsupervised data analysis clearly segregated the samples according to healthy controls and ZIKV patients as well as to convalescent and acute phases of infection along the first principal component (PC1) of the data. PC1 accounted for 42.5% of the variation in the measurements across the samples (Figure 2A). The second and third principal components (PC2, PC3), together accounted for 39.7% of the variation between measurements, and highlighted the differences between individuals. Along these axes, the patient samples from the convalescent phase of infection were closely grouped together compared to those from the acute phase of infection.

**Figure 2.**
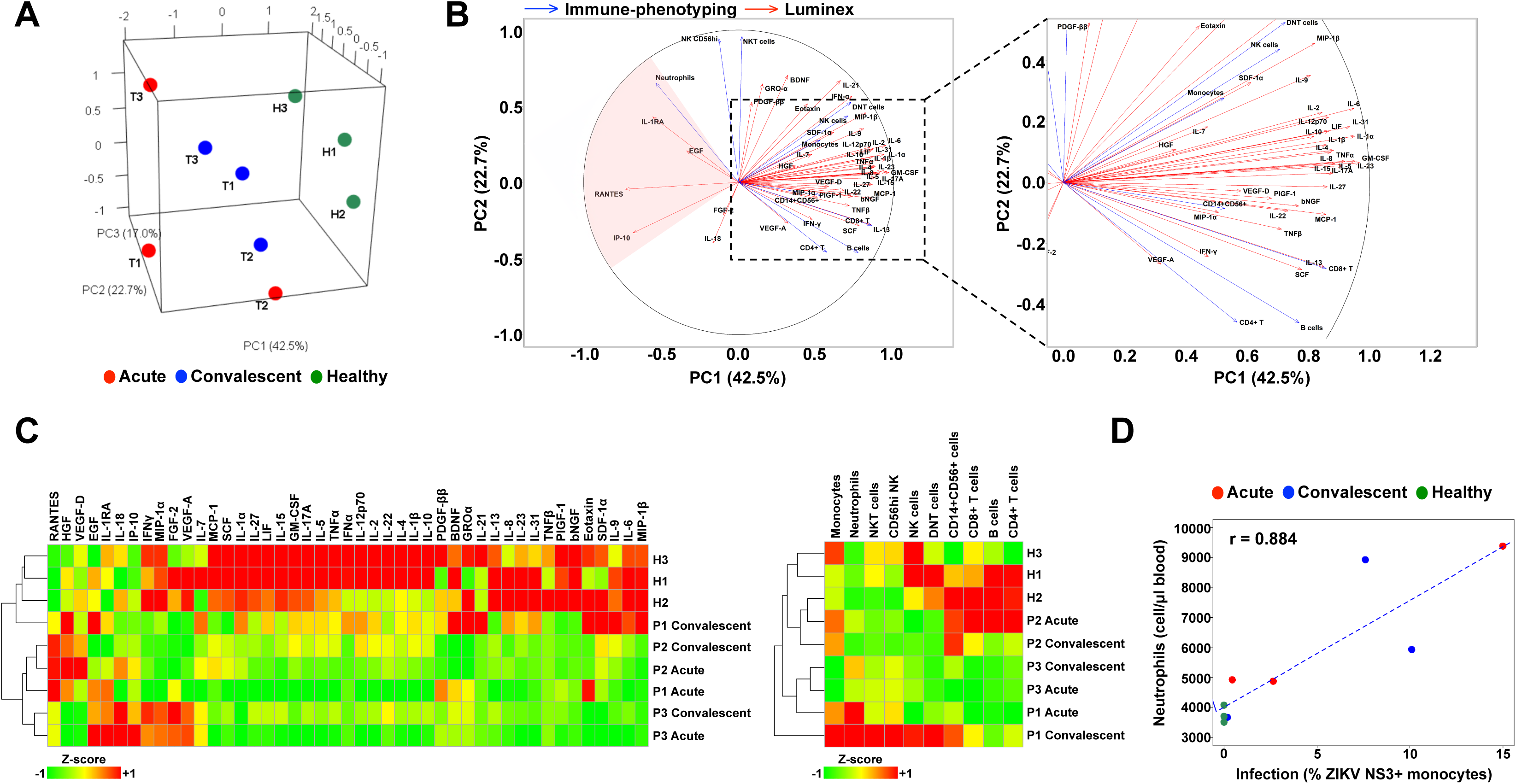
Immune characterization of ZIKV-infected pregnant women. Blood samples were obtained from three pregnant women during the acute and convalescent phases of ZIKV infection. The women were infected in different trimesters of their pregnancy: P1, trimester 1; P2, trimester 2; P3, trimester 3. Levels of peripheral blood immune cells and immune mediators were quantified by luminex and immunophenotyping, respectively. Healthy controls (n = 3) were included. (A) PCA scores of the infected and healthy women based on a collective analysis of immunophenotyping and cytokine measurements with Multiple Factor Analysis (MFA). (B) Correlation of immunophenotyping and cytokine measurements with the first two principal component axes (PC1, PC2) of the MFA. (C) Heatmaps comparing the cytokine and immunophenotyping data for each subject. (D) Correlation between infection (% ZIKV NS3+ monocytes) and neutrophil numbers in whole blood samples.

Correlation of measurements with PC1, which separated ZIKV patients and healthy controls, showed that a subset of immune mediators (IL-1RA, EGF, RANTES, IP-10) and neutrophils were associated with acute ZIKV infection (Figure 2B and 2C). Noteworthy, neutrophil numbers in the peripheral blood positively correlated with the percentage of ZIKV-infected (ZIKV NS3+) monocytes (Figure 2D and Table 1). Overall, the results obtained suggested that ZIKV infection during trimester 1 could lead to a more active pro-inflammatory response as observed in Patient 1.

### ZIKV infection during the first trimester of pregnancy triggered disparate eIF2 signaling and oxidative phosphorylation in the placenta

Transcriptome profiling of the placenta discs and fetal membranes of the placentas was used to investigate the cellular host response during ZIKV infection. Both tissues were digested to obtain a single-cell suspension: the placenta disc was further separated into CD45+ immune and CD45-non-immune fractions, to define the drivers of ZIKV-induced cellular changes. Tissue samples were collected in triplicate to account for tissue heterogeneity, except those obtained from Patient 3, where the sample was processed as a whole, due to technical difficulties. Placental samples from two healthy women were included as controls.

Over 517 million paired-end RNA-seq reads were obtained with a median of >15 million paired-end reads per sample mapped to the human transcriptome and 29,613 genes had a detectable expression level of >1 transcript per million (TPM) per sample. Principal component analysis (PCA) of the gene expression profiles segregated the samples by their origin (placenta disc or fetal membrane), regardless of whether they were derived from patients or controls (Figure 3A). Patient 1 was the only exception, as the gene expression data all clustered together (Figure 3A, cluster 4). This effect was further elaborated when PCA was performed separately for each tissue type (Figure 3B), where samples from Patient 1 segregated from the rest, indicating that the placenta from Patient 1 bore a distinct gene expression profile.

**Figure 3.**
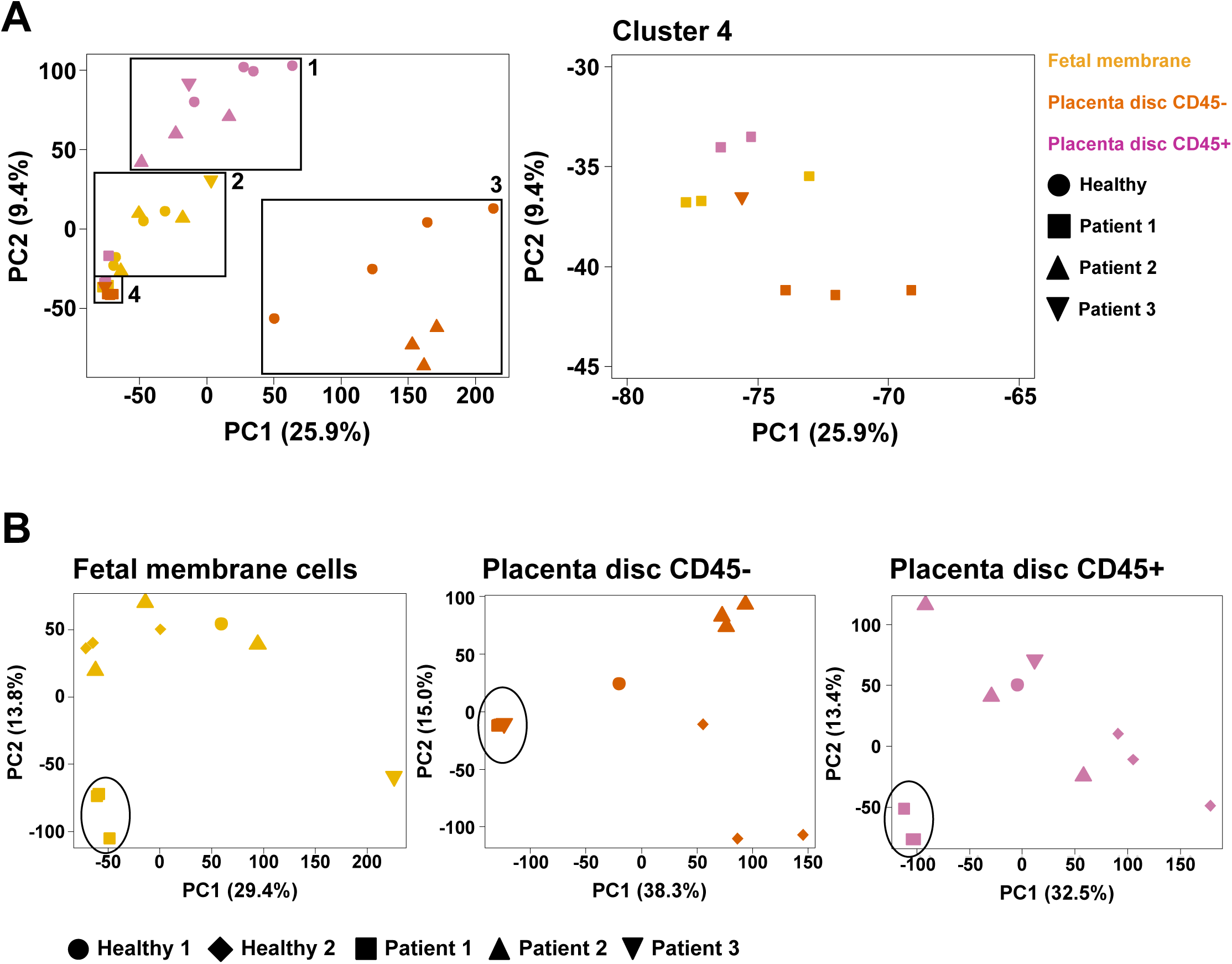
Principal component analysis of gene expression in placenta samples. Full-term placentas from ZIKV-infected pregnant patients were obtained and separated into the fetal membrane, and CD45-non-immune and CD45+ immune cells of the placenta disc. Placentas from two healthy women were obtained in parallel as a control. The transcriptomes of the three cell types (Placenta disc CD45+, Placenta disc Maternal CD45- and Fetal membrane cells) were assessed by RNA-sequencing (RNA-seq). PCA was performed by analyzing RNA-seq data either (A) collectively from all three cell types, or (B) individually as each specific cell type.

To determine the cellular differences in the placenta of Patient 1, the unique differential gene expression signature was compared to healthy controls. A large number of differentially expressed genes (DEGs) were identified between the placentas of Patient 1 and those from healthy controls (Placenta disc CD45+, n = 674; Placenta disc CD45-, n = 2,780; Fetal membrane, n = 2,467). Conversely, few to no DEGs were identified between the placentas derived from Patient 2 and healthy controls (Placenta disc CD45+, n = 0; Placenta disc CD45-, n = 51, Fetal membrane, n = 1) (Supplementary Figure 4A). Unfortunately, samples from Patient 3 could not be analyzed due to an insufficient number of replicates. Nevertheless, these findings in DEGs agree with the observations from PCA analysis, where samples from Patient 1 were distinct from the rest.

The potential functions of the DEGs in placental tissue from Patient 1 were analyzed using gene ontology (GO), pathway enrichment and gene set enrichment analysis (GSEA) to provide a global overview. In the CD45-fraction of the placenta disc from Patient 1, GO analysis identified several DEGs as ribonucleoproteins, mitochondrial genes and histones (Figure 4 and Supplementary Table 1). Ingenuity Pathway Analysis (IPA) identified the role of eIF2 signaling, as the main canonical pathway associated with these DEGs (Supplementary Figure 4B). The eIF2 pathway regulates global protein synthesis and translation in response to stress. Oxidative phosphorylation was also significantly modulated, where 90% of genes involved in this pathway showed a reduced expression by >2-fold compared to healthy controls. An increased abundance of histone transcripts during viral infection has been previously reported [26, 27] and was also observed in Patient 1 (Figure 4). The data also suggested potential modulation of oxidative phosphorylation and eIF2 signaling in both the CD45+ cells of the placenta disc and fetal membrane in the placenta from Patient 1 (Figure 4, Supplementary Figure 4B and Supplementary Table 2).

**Figure 4.**
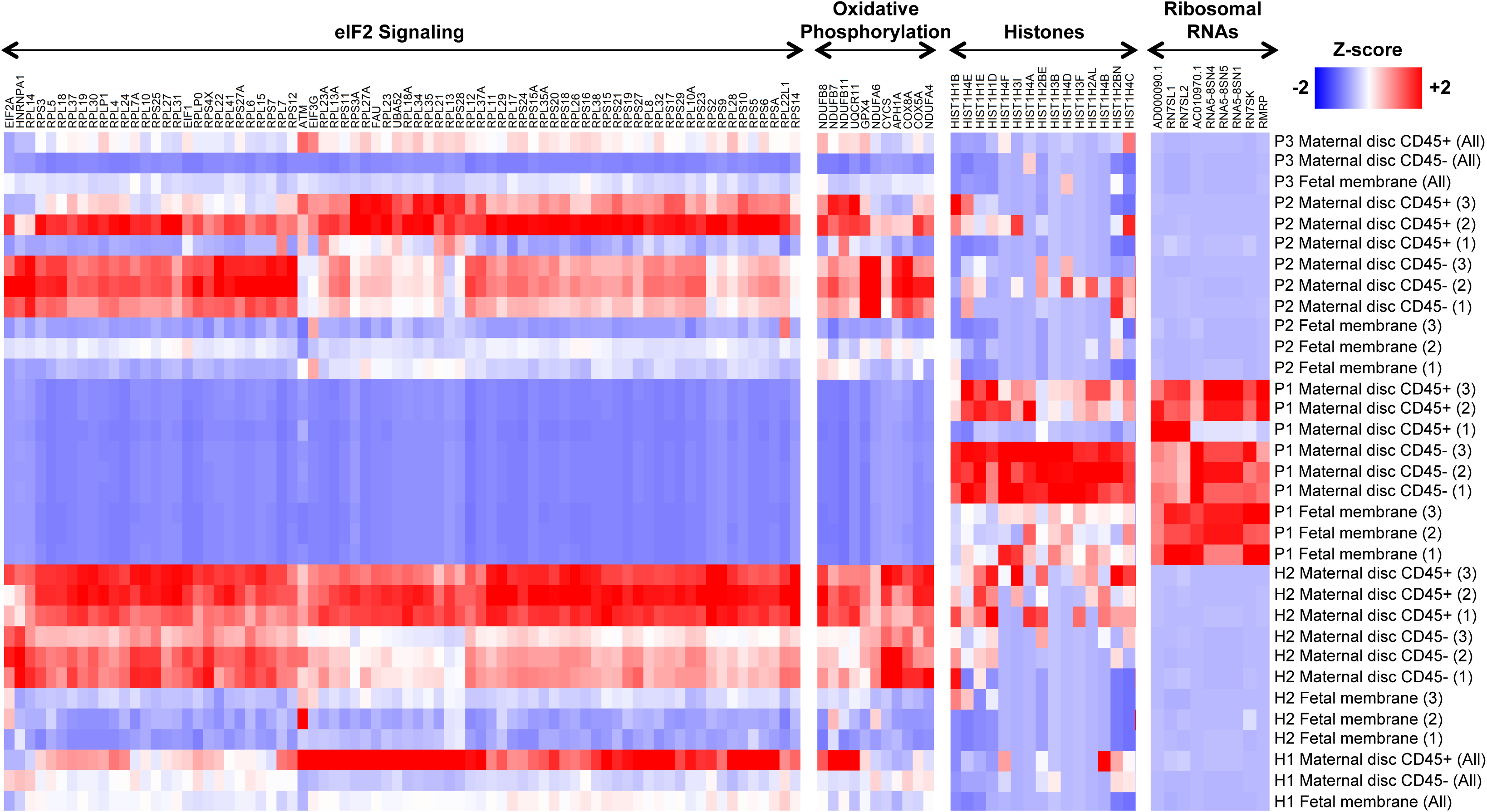
Placental gene expression associated with ZIKV infection. Heatmaps comparing gene expression in the different placental specimens (Placenta disc CD45+, Placenta disc CD45- and Fetal membrane cells) from ZIKV-infected women infected during the first (P1), second (P2) or third (P3) trimesters of pregnancy. Placental samples from two healthy women (H1, H2) were obtained in parallel as a control. Samples from H2, P1 and P2 were processed in triplicates. The most highly differentially regulated genes and pathways in P1 included EIF2 signaling, oxidative phosphorylation, histones and ribosomal RNAs. Gene expression measurements from RNA-seq in transcripts per million (TPM) scale were used to calculate a Z-score on each row.

To determine the percentages of various immune populations, immunophenotyping of the isolated CD45+ cells was performed (Figure 5A and Supplementary Figure 5). Correlating this immunophenotyping data with the first two principal components of the gene expression data (Figure 5B) corroborated that the increase in neutrophil numbers (Figure 5C) was clearly associated with gene expression changes observed in the placenta from Patient 1 compared to healthy placentas. In the fetal membrane of the placenta, several transcripts encoded by inflammatory response genes and pathways were increased in abundance, including NFKB1, IL1B, TNF, TLR2, TLR4, CXCL8, NLRP3 and NLRP6 (Supplementary Figure 4B and Supplementary Table 3). Taken together, these data suggest that ZIKV infection during the first trimester of pregnancy could have led to an upregulated abundance of transcripts targeting eIF2 signaling, oxidative phosphorylation and neutrophil numbers, as observed in Patient 1.

**Figure 5.**
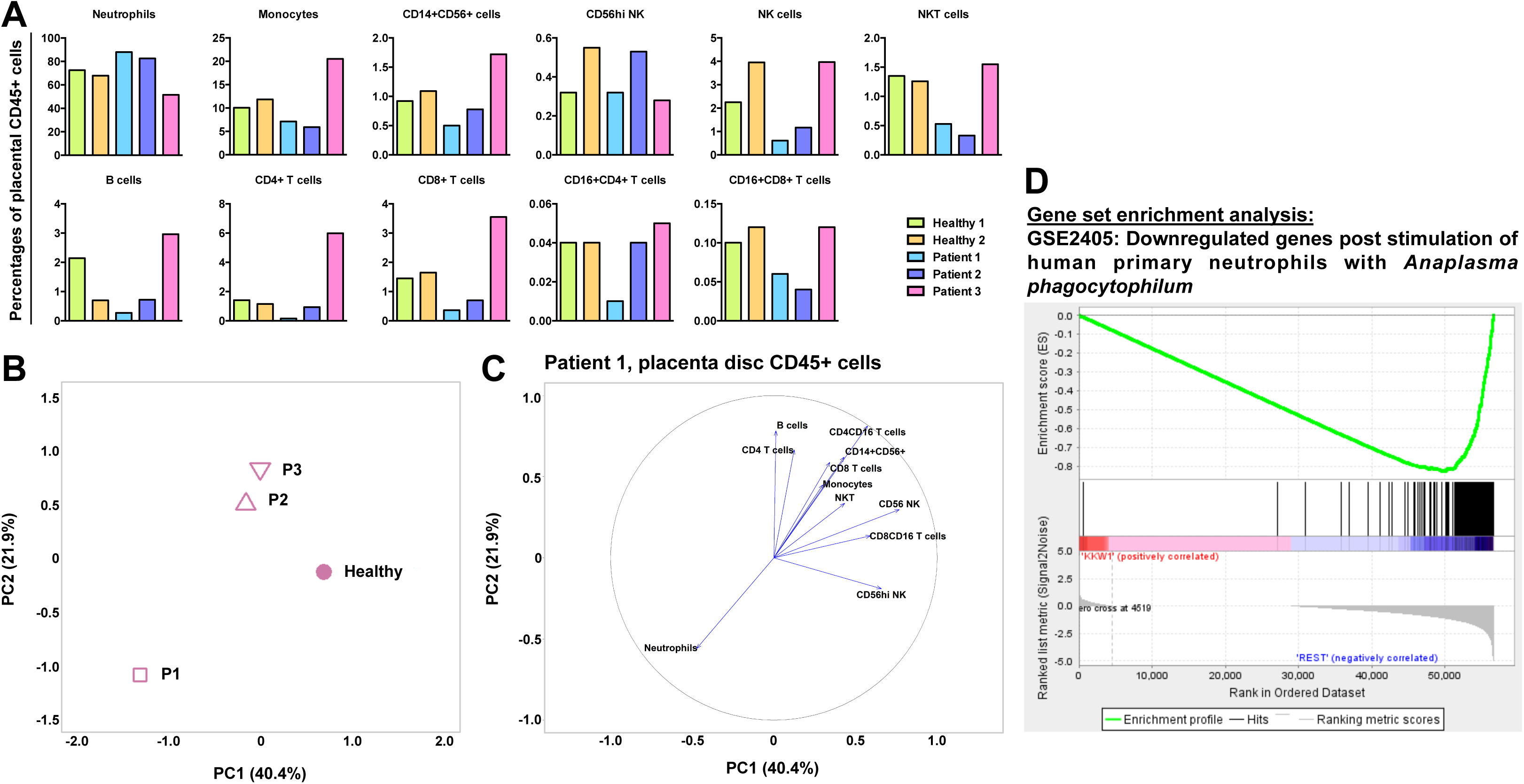
ZIKV infection during pregnancy first trimester triggers disparate cellular response in placenta disc CD45+ cells. (A) Relative proportions of immune subsets in placenta disc CD45+ cells in ZIKV-infected patients (n=3) and healthy controls (n=2). (B) PCA of gene expression profiles of placenta disc CD45+ cells from ZIKV patients and healthy controls. The PCA was performed on 30,717 genes selected by standard deviation > 0. (C) Correlation plot of immune-cell subset frequencies in placenta disc CD45+ cells with the first two principal components (PC1, PC2) of the gene expression profiles. (D) Enrichment plot of the gene set GSE2405_0H_VS_9H_A_PHAGOCYTOPHILUM_STIM_NEUTROPHIL_DN, which was reported by GSEA as most enriched among all immunological gene sets (C7 of MSigDB) in the placenta disc CD45+ fraction of Patient 1 vs. Healthy placenta. The profile shows the running enrichment score (green curve) and the positions of gene-set members (black vertical bars) on the rank-ordered list of differential gene expression comparing the Patient 1 vs. Healthy placenta disc CD45+ fraction.

## Discussion

The emergence of ZIKV in recent years came as a surprise and the apparent association of ZIKV infection with neurological complications in developing fetuses, namely congenital microcephaly, further alarmed the world. ZIKV is capable of infecting neural cells [28–33], providing a direct causative role of the virus in causing neurological impairment. In animal studies, ZIKV infection of immuno-compromised pregnant female mice resulted in premature fetal death [34]; ZIKV was found in the placenta in these mice, leading to active placental infection, apoptosis and vascular damage [34]. In humans, it is worth noting that the estimated prevalence of microcephaly in babies born to ZIKV-infected pregnant women was ∼2.3% [35]. This estimate, however, considers infections that have occurred in any of the three trimesters [35]. The likelihood of microcephaly is thought to be higher if ZIKV infection occurs during the first trimester [11].

This study recruited three pregnant women who were infected with ZIKV during different trimesters of pregnancy. Despite the low number of patients recruited (one patient per trimester), this remains an excellent opportunity to obtain an insight into the impact of ZIKV infection during different trimesters on host cellular responses. Nevertheless, a series of histologic, immunologic and transcriptomic analyses were used to determine the impact of ZIKV infection on pregnancy progression and the cellular responses of the host towards the infection. All three babies were born at term with no observable birth defects, and all of the ZIKV-infected mothers were clinically well, as with majority of ZIKV patients [14]. However, different degrees of infection were apparent in the level of CD14+ monocytes between the three patients. This difference led to a disparate immune response exhibited by the patients. As confirmed by MFA, several immune mediators were associated with the acute phase of ZIKV infection, including IL-1RA, EGF, RANTES and IP-10 [14]. Interestingly, EGF and IP-10 are typically expressed at higher levels in women carrying a fetus with developmental anomalies [36]. The association of neutrophils with the acute phase should also be given more attention since most studies have focused on the role of monocytes during ZIKV infection thus far [37, 38]. Neutrophils have been shown to infiltrate the central nervous system of ZIKV-infected immuno-compromised mice [39], and may therefore be a causative element in Zika-induced neurological damage.

ZIKV NS3 antigen was detected in placental tissues of all three patients at full term, indicating a strong tropism of the virus to the placenta regardless of the trimester of infection. The ZIKV NS3 antigen was detected in Hofbauer cells, placental macrophages [9], but not in trophoblasts. The susceptibility of trophoblastic cells to ZIKV infection is still uncertain, with reports indicating either permissive or resistance towards ZIKV infection [9, 40, 41]. Interestingly, the localization of the ZIKV NS3 antigen to CD163+ Hofbauer cells did not corroborate with positive detection of ZIKV RNA load in the placental tissue. This finding may indicate that the virus has been cleared and the NS3 antigens are merely phagocytosed, non-infectious viral remnants. This observation was further supported by the lack of any pathological insults (villitis, intervillositis, chorioamnionitis, or funisitis) to the placental tissues in any of the three patients as well as the lack of detectable viral RNA by RNAseq, performed at the resolution sequenced. This observation also indicated an absence of overt inflammation, which permitted the pregnancy to progress to term without complications. This result was further confirmed by the absence of Hofbauer cell-mediated hyperplasia in the chorionic villi, which has been previously associated with progression of congenital ZIKV infection [41].

The transcriptomic profiling of placental cells showed that the cellular responses in Patient 1 (infected during the first trimester) were quite different from the other two patients and healthy controls for all three cell types analyzed: placenta disc CD45+, placenta disc CD45- and fetal membrane cells. GSEA revealed that a subset of genes in the CD45+ cells of Patient 1 (Figure 5D) resembled those documented in the bacterium *Anaplasma phagocytophilum*-stimulated human primary neutrophils [42]. This finding is not surprising, given that isolated CD45+ cells comprise a high proportion of neutrophils and thus would contribute the most to the transcriptome. More importantly, this finding also indicates the vast number of neutrophils present in the placenta during pregnancy. Their participation should not be taken lightly as this cell type may be involved in infertility, preeclampsia and even fetal loss. Neutrophils also participate in implantation, angiogenesis, spiral artery modification and parturition [43].

The DEGs in the placenta disc, both CD45+ and CD45-fractions, and fetal membrane of Patient 1 were strongly associated with eIF2 signaling, oxidative phosphorylation and mitochondrial dysfunction. eIF2 signaling is crucial in regulating translational initiation leading to protein synthesis, and participates in cellular stress responses, erythropoiesis and immune responses to viral infection [44]. Perturbed eIF2 signaling in placental development has been associated with intrauterine growth restriction of developing fetuses [45]. In rats, eIF2 signaling was also shown to be heavily involved in the regulation of placental and fetal development [46]. Likewise, oxidative phosphorylation and mitochondrial dysfunction have been associated with a range of gestational disorders [47].

These findings described herein can be translated into the clinical prenatal screening procedures in ZIKV-infected or any arbovirus-infected pregnant patients. In such circumstances, abnormalities in the numbers of leukocytes [48, 49] as well as biomarkers indicative of ongoing inflammation [50] or oxidative stress [51] and presence of any microbial infections [52] should be screened as advised by the clinic. Likewise, RNA from the amniotic fluid or amniocytes, can be profiled to determine the transcript abundance of molecules involved in important biological pathways [53]. Importantly, fetal development (e.g. blood flow, heart rate, head circumference) should be closely monitored by ultrasound scanning [54] to detect for unwanted anomalies that may surface in the later part of life [25]. Adopting these steps will potentially improve the clinical care administered to affected patients, by identifying potential danger signs associated with ZIKV infection.

## Acknowledgements

We would like to thank Ivy Low, Seri Mustafah, and Anis Larbi (SIgN flow cytometry team) and the SIgN Immunomonitoring Group, for their assistance. We are also grateful to Fatima Yturriaga and Emmerie Wong Phaik Yen of KKH for the patient recruitment, Kai-Er Eng and Guillaume Carissimo of SIgN and Jessica Tamanini of Insight Editing London for editing the manuscript prior to submission. We would also like to thank all study participants for their participation in this study.

## References

1. Korzeniewski, K., D. Juszczak, and E. Zwolinska, Zika - another threat on the epidemiological map of the world. Int Marit Health, 2016. 67(1): p. 31–7.

2. Brasil, P., et al., Zika Virus Infection in Pregnant Women in Rio de Janeiro - Preliminary Report. N Engl J Med, 2016.

3. Mlakar, J., et al., Zika Virus Associated with Microcephaly. N Engl J Med, 2016. 374(10): p. 951–8.

4. Mor, G. and I. Cardenas, The immune system in pregnancy: a unique complexity. Am J Reprod Immunol, 2010. 63(6): p. 425–33.

5. Warning, J.C., S.A. McCracken, and J.M. Morris, A balancing act: mechanisms by which the fetus avoids rejection by the maternal immune system. Reproduction, 2011. 141(6): p. 715–24.

6. Sappenfield, E., D.J. Jamieson, and A.P. Kourtis, Pregnancy and susceptibility to infectious diseases. Infect Dis Obstet Gynecol, 2013. 2013: p. 752852.

7. Burton, G.J. and E. Jauniaux, What is the placenta? Am J Obstet Gynecol, 2015. 213(4 Suppl): p. S6 e1, S6-8.

8. Robbins, J.R. and A.I. Bakardjiev, Pathogens and the placental fortress. Curr Opin Microbiol, 2012. 15(1): p. 36–43.

9. Quicke, K.M., et al., Zika Virus Infects Human Placental Macrophages. Cell Host Microbe, 2016. 20(1): p. 83–90.

10. Eppes, C., et al., Testing for Zika virus infection in pregnancy: key concepts to deal with an emerging epidemic. Am J Obstet Gynecol, 2017. 216(3): p. 209–225.

11. Johansson, M.A., et al., Zika and the Risk of Microcephaly. N Engl J Med, 2016. 375(1): p. 1–4.

12. Kam, Y.W., et al., Cross-reactive dengue human monoclonal antibody prevents severe pathologies and death from Zika virus infections. JCI Insight, 2017. 2(8).

13. Lum, F.M., et al., Sensitive detection of Zika virus antigen in patients’ whole blood as an alternative diagnostic approach. J Infect Dis, 2017. 216(2): p. 182–190.

14. Lum, F.M., et al., Longitudinal study of cellular and systemic cytokines signatures define the dynamics of a balanced immune environment in disease manifestation in Zika virus-infected patients. J Infect Dis, 2018. 218(5): p. 814–824.

15. Martin, M., Cutadapt removes adapter sequences from high-throughput sequencing reads. EMBnet. journal, 2011. 17(1): p. pp-10.

16. Harrow, J., et al., GENCODE: the reference human genome annotation for The ENCODE Project. Genome Res, 2012. 22(9): p. 1760–74.

17. Patro, R., et al., Salmon provides fast and bias-aware quantification of transcript expression. Nat Methods, 2017. 14(4): p. 417–419.

18. Soneson, C., M.I. Love, and M.D. Robinson, Differential analyses for RNA-seq: transcript-level estimates improve gene-level inferences. F1000Res, 2015. 4: p. 1521.

19. Lê, S., J. Josse, and F. Husson, FactoMineR: an R package for multivariate analysis. Journal of statistical software, 2008. 25(1): p. 18.

20. Lun, A.T., Y. Chen, and G.K. Smyth, It’s DE-licious: A Recipe for Differential Expression Analyses of RNA-seq Experiments Using Quasi-Likelihood Methods in edgeR. Methods Mol Biol, 2016. 1418: p. 391–416.

21. Huang da, W., B.T. Sherman, and R.A. Lempicki, Systematic and integrative analysis of large gene lists using DAVID bioinformatics resources. Nat Protoc, 2009. 4(1): p. 44–57.

22. Subramanian, A., et al., Gene set enrichment analysis: a knowledge-based approach for interpreting genome-wide expression profiles. Proc Natl Acad Sci U S A, 2005. 102(43): p. 15545–50.

23. Escofier, B. and J. Pagès, Multiple factor analysis (AFMULT package). Comput Stat Data Anal, 1994. 18(1): p. 121–140.

24. Svensson, J., et al., Macrophages at the fetal-maternal interface express markers of alternative activation and are induced by M-CSF and IL-10. J Immunol, 2011. 187(7): p. 3671–82.

25. Nielsen-Saines, K., et al., Delayed childhood neurodevelopment and neurosensory alterations in the second year of life in a prospective cohort of ZIKV-exposed children. Nat Med, 2019.

26. Guerra, S., et al., Microarray analysis reveals characteristic changes of host cell gene expression in response to attenuated modified vaccinia virus Ankara infection of human HeLa cells. J Virol, 2004. 78(11): p. 5820–34.

27. Hoover, S.E., et al., Downregulation of varicella-zoster virus (VZV) immediate-early ORF62 transcription by VZV ORF63 correlates with virus replication in vitro and with latency. J Virol, 2006. 80(7): p. 3459–68.

28. Lanko, K., et al., Replication of the Zika virus in different iPSC-derived neuronal cells and implications to assess efficacy of antivirals. Antiviral Res, 2017. 145: p. 82–86.

29. Lum, F.M., et al., Zika Virus Infects Human Fetal Brain Microglia and Induces Inflammation. Clin Infect Dis, 2017. 64(7): p. 914–920.

30. McGrath, E.L., et al., Differential Responses of Human Fetal Brain Neural Stem Cells to Zika Virus Infection. Stem Cell Reports, 2017. 8(3): p. 715–727.

31. Olmo, I.G., et al., Zika Virus Promotes Neuronal Cell Death in a Non-Cell Autonomous Manner by Triggering the Release of Neurotoxic Factors. Front Immunol, 2017. 8: p. 1016.

32. Simonin, Y., et al., Zika Virus Strains Potentially Display Different Infectious Profiles in Human Neural Cells. EBioMedicine, 2016. 12: p. 161–169.

33. Tang, H., et al., Zika Virus Infects Human Cortical Neural Progenitors and Attenuates Their Growth. Cell Stem Cell, 2016. 18(5): p. 587–90.

34. Miner, J.J., et al., Zika Virus Infection during Pregnancy in Mice Causes Placental Damage and Fetal Demise. Cell, 2016. 165(5): p. 1081–91.

35. Coelho, A.V.C. and S. Crovella, Microcephaly Prevalence in Infants Born to Zika Virus-Infected Women: A Systematic Review and Meta-Analysis. Int J Mol Sci, 2017. 18(8).

36. Kam, Y.W., et al., Specific Biomarkers Associated With Neurological Complications and Congenital Central Nervous System Abnormalities From Zika Virus-Infected Patients in Brazil. J Infect Dis, 2017. 216(2): p. 172–181.

37. Foo, S.S., et al., Asian Zika virus strains target CD14+ blood monocytes and induce M2-skewed immunosuppression during pregnancy. Nat Microbiol, 2017.

38. Lum, F.M., et al., Zika virus infection preferentially counterbalances human peripheral monocyte and/or NK-cell activity. bioRxiv, 2017.

39. Manangeeswaran, M., D.D. Ireland, and D. Verthelyi, Zika (PRVABC59) Infection Is Associated with T cell Infiltration and Neurodegeneration in CNS of Immunocompetent Neonatal C57Bl/6 Mice. PLoS Pathog, 2016. 12(11): p. e1006004.

40. Bayer, A., et al., Type III Interferons Produced by Human Placental Trophoblasts Confer Protection against Zika Virus Infection. Cell Host Microbe, 2016. 19(5): p. 705–12.

41. Schwartz, D.A., Viral infection, proliferation, and hyperplasia of Hofbauer cells and absence of inflammation characterize the placental pathology of fetuses with congenital Zika virus infection. Arch Gynecol Obstet, 2017. 295(6): p. 1361–1368.

42. Borjesson, D.L., et al., Insights into pathogen immune evasion mechanisms: Anaplasma phagocytophilum fails to induce an apoptosis differentiation program in human neutrophils. J Immunol, 2005. 174(10): p. 6364–72.

43. Giaglis, S., et al., Neutrophil migration into the placenta: Good, bad or deadly? Cell Adh Migr, 2016. 10(1-2): p. 208–25.

44. Wek, R.C., Role of eIF2alpha Kinases in Translational Control and Adaptation to Cellular Stress. Cold Spring Harb Perspect Biol, 2018. 10(7).

45. Yung, H.W., et al., Evidence of placental translation inhibition and endoplasmic reticulum stress in the etiology of human intrauterine growth restriction. Am J Pathol, 2008. 173(2): p. 451–62.

46. Gaccioli, F., et al., Maternal overweight induced by a diet with high content of saturated fat activates placental mTOR and eIF2alpha signaling and increases fetal growth in rats. Biol Reprod, 2013. 89(4): p. 96.

47. Holland, O., et al., Review: Placental mitochondrial function and structure in gestational disorders. Placenta, 2017. 54: p. 2–9.

48. Park, C.W., et al., The frequency and clinical significance of intra-amniotic inflammation defined as an elevated amniotic fluid matrix metalloproteinase-8 in patients with preterm labor and low amniotic fluid white blood cell counts. Obstet Gynecol Sci, 2013. 56(3): p. 167–75.

49. Romero, R., et al., Amniotic fluid white blood cell count: a rapid and simple test to diagnose microbial invasion of the amniotic cavity and predict preterm delivery. Am J Obstet Gynecol, 1991. 165(4 Pt 1): p. 821–30.

50. Abdallah, M.W., et al., Amniotic fluid inflammatory cytokines: potential markers of immunologic dysfunction in autism spectrum disorders. World J Biol Psychiatry, 2013. 14(7): p. 528–38.

51. Kacerovsky, M., et al., Amniotic fluid markers of oxidative stress in pregnancies complicated by preterm prelabor rupture of membranes. J Matern Fetal Neonatal Med, 2014: p. 1–10.

52. Hayat, M., et al., A very unusual complication of amniocentesis. Clin Case Rep, 2015. 3(6): p. 345–8.

53. Kang, J.H., et al., Comparative Transcriptome Analysis of Cell-Free Fetal RNA from Amniotic Fluid and RNA from Amniocytes in Uncomplicated Pregnancies. PLoS One, 2015. 10(7): p. e0132955.

54. Rayburn, W.F., J.A. Jolley, and L.L. Simpson, Advances in ultrasound imaging for congenital malformations during early gestation. Birth Defects Res A Clin Mol Teratol, 2015. 103(4): p. 260–8.

