## Supplementary Figure 1 for "Trimester-specific Zika virus infection affects placental responses in women"

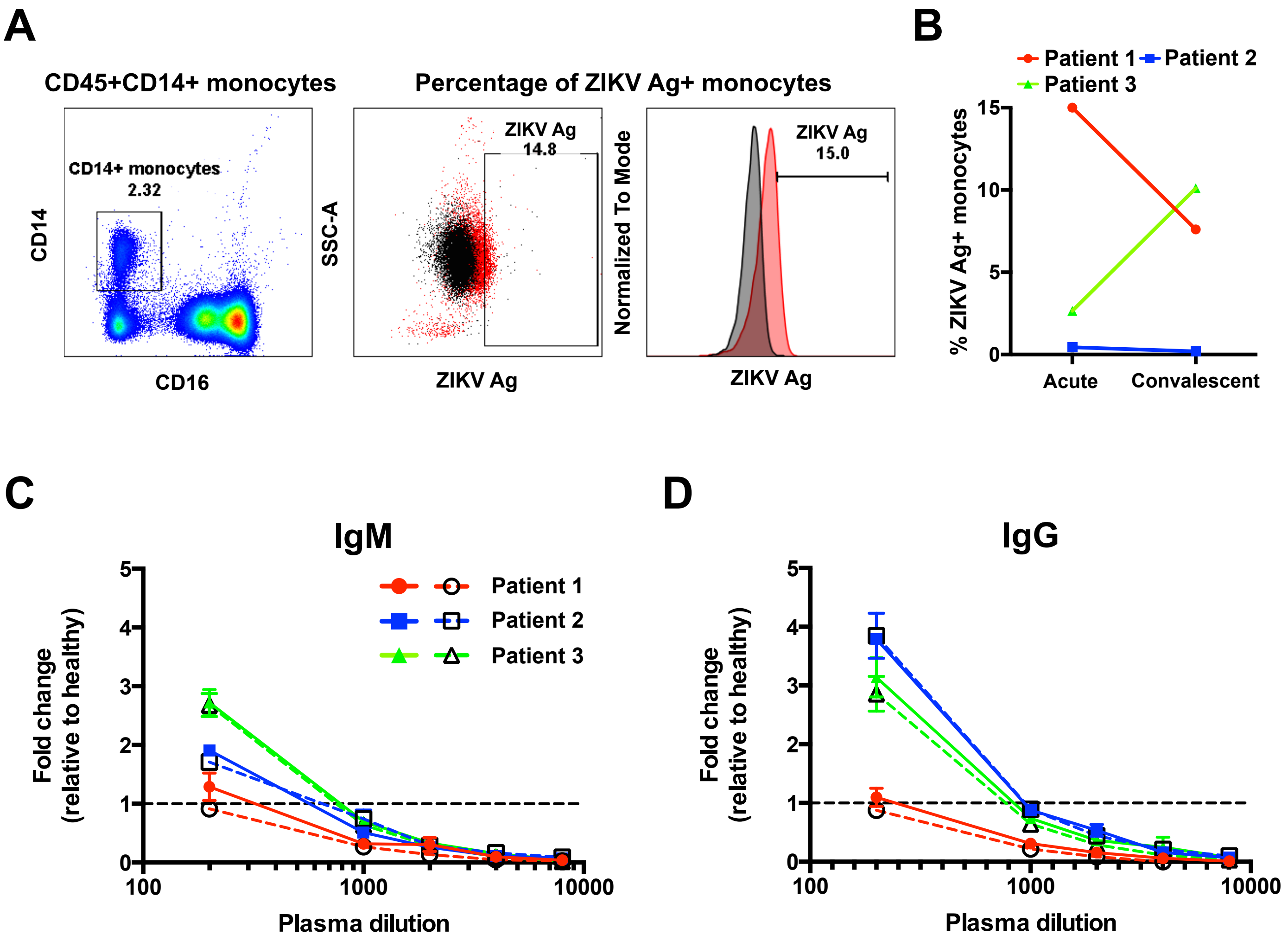


**Supplementary Figure 1. Detection of positive ZIKV infection.** Three pregnant women infected with ZIKV in different trimesters of pregnancy were recruited. (A) Representative plots from Patient 1 showing the detection of ZIKV antigen in blood CD45+CD14+ monocytes by flow cytometry. (B) Assessment of ZIKV infection in pregnant patients presented as paired-line graph to illustrate the transition of infection levels in each patient from the acute through to the convalescent phase. The levels of ZIKV-specific IgM (C) and IgG (D) antibodies present in the plasma obtained during acute (colored symbol) and convalescent (clear symbol) disease phases were titered with a ZIKV-virion based ELISA. Samples were tested in quadruplicate and are presented as the signal fold change relative to the healthy samples (normalized to 1). Continuous and broken lines represent samples from acute and convalescent phases, respectively.

**Supplementary Figure 1 Lum et al., 2019**
