## Supplementary Figure 2 for "Trimester-specific Zika virus infection affects placental responses in women"

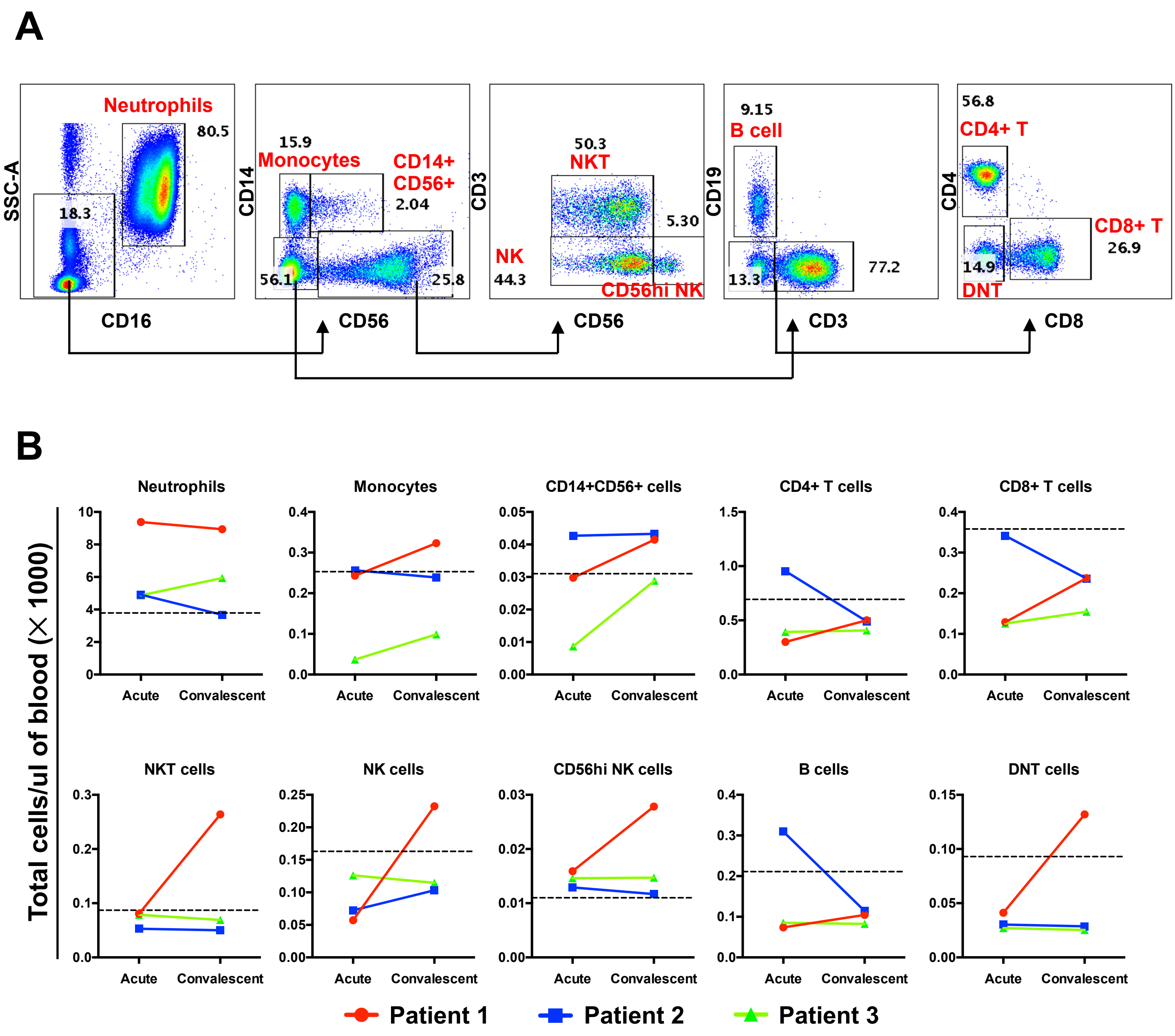


**Supplementary Figure 2. Peripheral blood immune-cell quantification.** Cellular numbers of peripheral blood immune cells were determined by immunophenotyping with a panel of CD45, CD14, CD3, CD4, CD8, CD19, CD16 and CD56 antibodies. (A) Gating strategy for immunophenotyping. The scatter plots show a representative donor. (B) Cellular numbers for each peripheral blood immune-cell subset were calculated with the following formula: [Percentages of specific immune subset × total leucocyte numbers = cellular numbers of specific immune subset]. Total leucocyte numbers were obtained using a hematology analyzer. Data are plotted in paired-line graphs to illustrate the changes in cellular numbers transiting between the acute and convalescent phases of ZIKV infection. The horizontal dotted line in each plot indicates the mean quantity for each specific immune subset, derived from a group of 14 healthy adults.

**Supplementary Figure 2 Lum et al., 2019**
