## Supplementary Figure 3 for "Trimester-specific Zika virus infection affects placental responses in women"

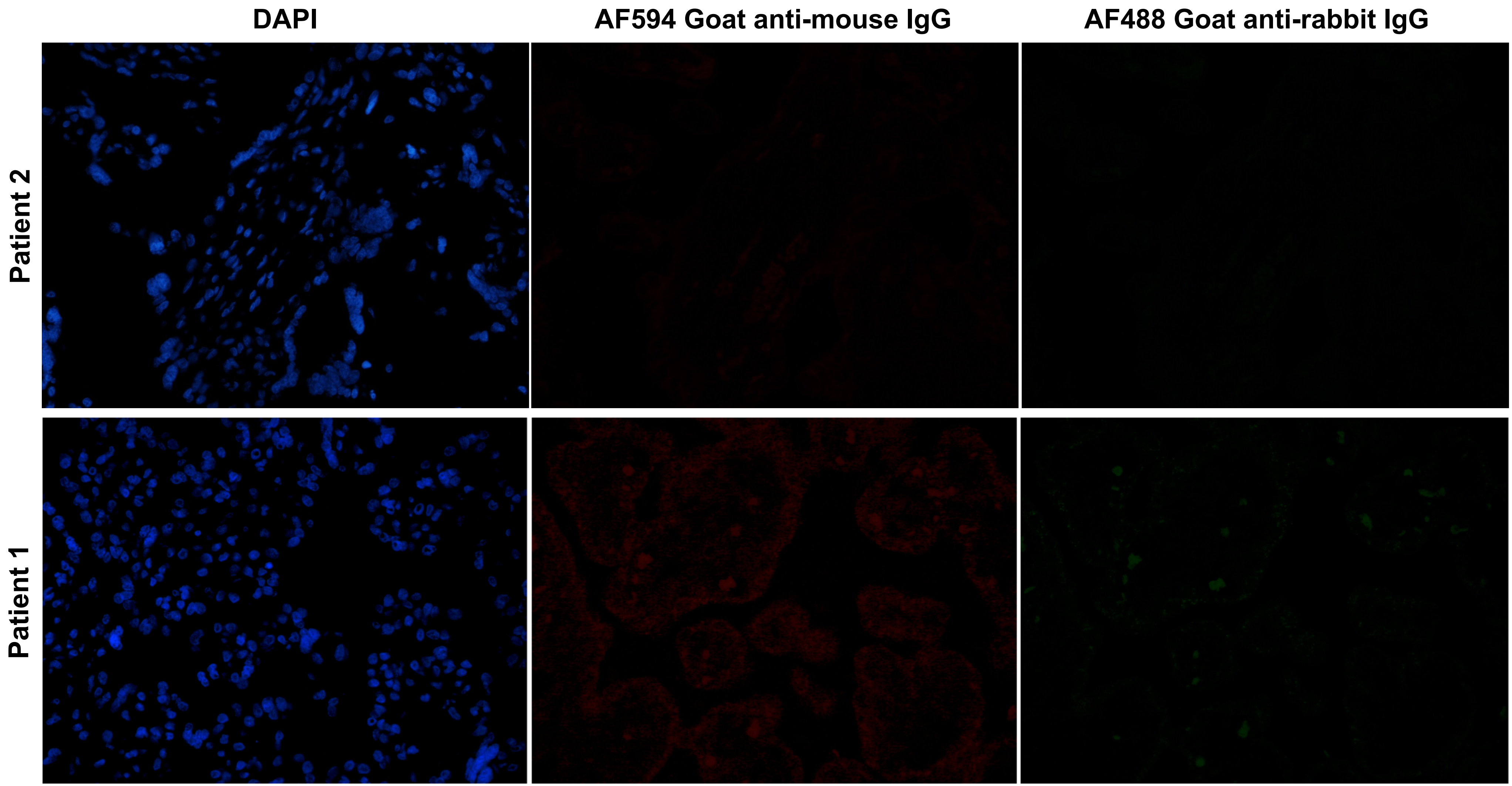


**Supplementary Figure 3. Secondary antibody staining of placenta discs.** Placenta discs of both patient 1 and patient 2 were stained with only secondary antibody AF594 Goat anti-mouse IgG and AF488 Goat anti-rabbit IgG. Images were captured at 40X magnification.

**Supplementary Figure 3 Lum et al., 2019**
