## Supplementary Figure 4 for "Trimester-specific Zika virus infection affects placental responses in women"

**A**

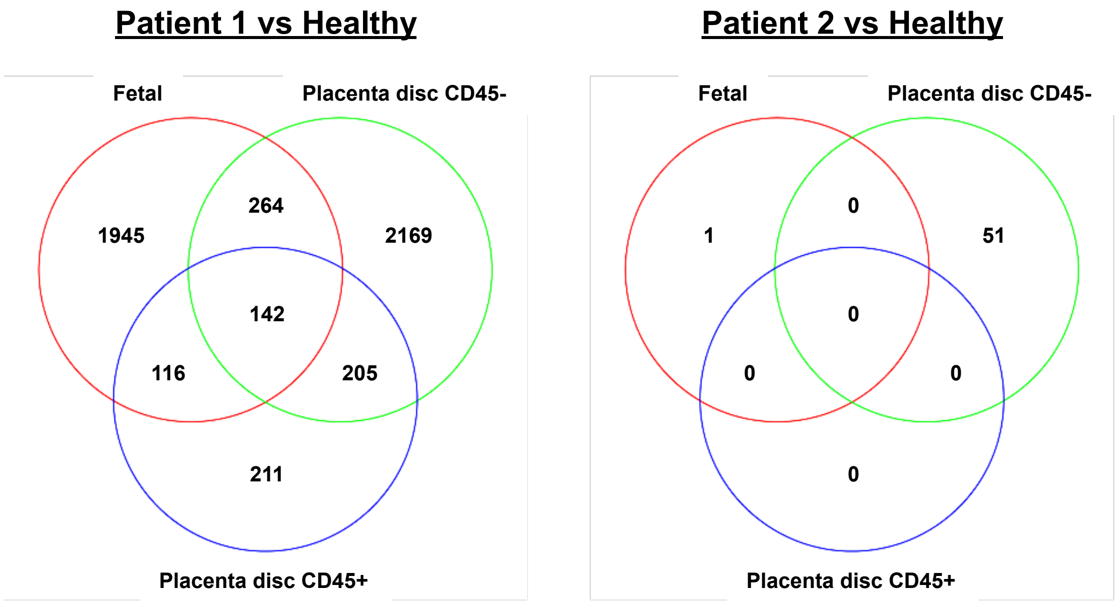


**B**

| Placenta disc CD45+ cells | P-values |
|---|---|
| ***Oxidative phosphorylation*** | 8.93E-25 |
| ***Mitochondrial dysfunction*** | 9.15E-21 |
| ***EIF2 signaling*** | 3.63E-12 |
| Sirtuin signaling pathway | 3.70E-10 |
| ***Regulation of eIF4 and p70S6K signaling*** | 1.00E-05 |
| Systemic Lupus Erythrematosis signaling | 4.55E-05 |
| Estrogen receptor signaling | 1.24E-04 |
| mTOR signaling | 5.28E-04 |
| Assembly of RNA polymerase II complex | 7.63E-04 |
| Pathogenesis of multiple sclerosis | 8.01E-04 |

| Placenta disc CD45- cells | P-values |
|---|---|
| ***EIF2 signaling*** | 5.19E-74 |
| ***Regulation of eIF4 and P70S6K signaling*** | 4.77E-24 |
| mTOR signaling | 1.07E-22 |
| Antigen presentation pathway | 2.67E-08 |
| Communication between innate and adaptive | 1.70E-06 |
| ***Mitochondrial dysfunction*** | 2.20E-05 |
| Autoimmune thyroid disease signaling | 2.96E-05 |
| Allograft rejection signaling | 3.10E-05 |
| ***Oxidative phosphorylation*** | 3.90E-05 |
| B cell development | 5.10E-05 |

| Fetal membrane cells | P-values |
|---|---|
| ***EIF2 Signaling*** | 1.52E-09 |
| ***Oxidative phosphorylation*** | 3.31E-08 |
| Neuroinflammation signaling pathway | 5.58E-07 |
| Sirtuin signaling pathway | 8.91E-06 |
| ***Regulation of eIF4 and P70S6K signaling*** | 1.28E-05 |
| ***Mitochondrial dysfunction*** | 1.38E-05 |
| TREM1 signaling | 1.58E-05 |
| IL-10 signaling | 3.22E-05 |
| IL-6 signaling | 3.26E-05 |
| Role of JAK family kinases in IL-6-type- | 6.23E-05 |

**Strength of association**

Low — High

**Supplementary Figure 4. Differentially expressed genes (DEGs) in placental specimens.** Full-term placentas from ZIKV-infected pregnant patients (patient 1 and 2; infected during the first and second pregnancy trimester respectively) were obtained and specific fetal membrane cells, placenta disc CD45- and CD45+ cells were isolated. Placentas from two healthy women were obtained in parallel as a control. The transcriptomes of the three cell types were subsequently assessed by RNA-sequencing. (A) Numbers of unique and common DEGs obtained between the various samples are depicted. (B) IPA analysis of DEGs contrasting to the healthy controls in the placenta disc (CD45+ and CD45-) and fetal membrane cells. The top 10 canonical pathways associated with each cell type are shown. Common canonical pathways are indicated in bold italics.

**Supplementary Figure 4 Lum et al., 2019**
