## Supplementary Figure 5 for "Trimester-specific Zika virus infection affects placental responses in women"

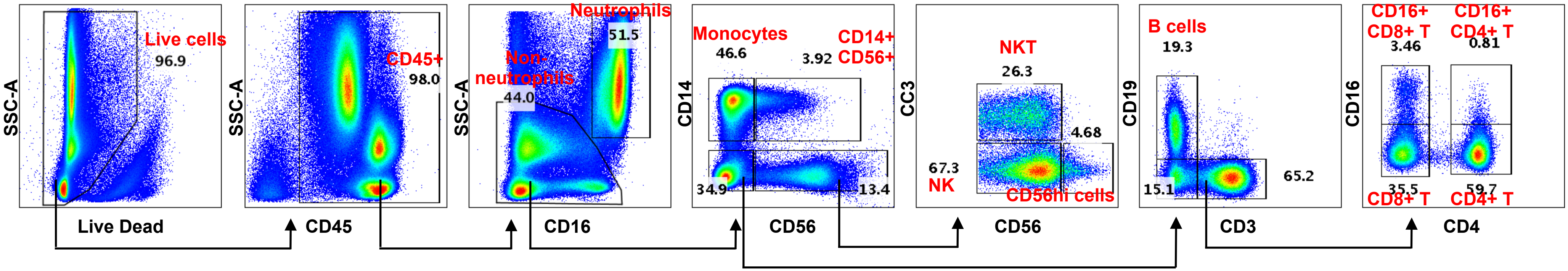


**Supplementary Figure 5. Immunophenotyping of full-term placentas.** Percentages of specific immune subsets were determined by immunophenotyping using a panel of CD45, CD14, CD3, CD4, CD19, CD16 and CD56 antibodies. The gating strategy for immunophenotyping is illustrated from a representative donor (placenta disc).

**Supplementary Figure 5 Lum et al., 2019**
